# Physiological barriers for glucose utilization in *Methanosarcina acetivorans*

**DOI:** 10.1101/2024.08.21.609014

**Authors:** Christian Sattler, Marcus Richter, Michael Rother

## Abstract

Methanogenesis is a key aspect of anaerobic biomass degradation, and thus of global importance. While methanogenic archaea (methanogens) are ubiquitous in anaerobic habitats, the range of substrates they utilize is very limited. Most methanogens are able to grow chemolithotrophically with H_2_+CO_2_, some clades can utilize methylated compounds and/or acetate as well. Organotrophic compounds like amino acids, lipids, nucleotides, or carbohydrates, are not known to support growth of methanogens. This inability to utilize “upstream” intermediates of anaerobic biomass degradation is remarkable considering the presence of cellular metabolism that such intermediates could feed into. Here, we addressed the question why the model methanogen *Methanosarcina acetivorans*, despite its gluconeogenic and glycolytic capacity, is unable to utilize glucose for methanogenesis and growth. Complementing heterologously with a glucose uptake facilitator allowed a recombinant *M. acetivorans* strain to convert glucose to methane at a low rate. However, growth with glucose was not observed, neither as energy source nor as carbon source. Instead, methylotrophic growth of the transgenic strain was impaired in a glucose-dependent fashion, which was aggravated when also glucokinase was heterologuously produced. Glucose-dependent growth inhibition coincided with a significant – and microscopically visible – accumulation of intracellular carbohydrate. Since the glucose-utilizing trait is rapidly lost during methylotrophic growth, accumulating growth-inhibiting metabolites probably makes methanogenic and glycolytic catabolism incompatible. Thus, extensive efforts in strain development would be required to enable direct glucose utilization for methanogenesis.

**Importance:** The known range of methanogenic growth substrates is very limited and the most reduced carbon compound to support organotrophic growth of *M. acetivorans* is pyruvate. Yet, gluconeogenesis and glycolysis, i. e., mobilization of stored glycogen, which is known to occur in *M. acetivorans* makes the capacity of glycolytic methanogenesis in this organism seem feasible. Although this trait could be engineered in *M. acetivorans* by introducing a single gene, the finding that it could not sustain growth of the organism, not even as an anabolic carbon source, demonstrates that methanogenesis is not easily compatible with carbohydrate utilization, thus probably requiring extensive metabolic rewiring.

## Introduction

Phototrophic (or chemotrophic) synthesis of biomass and its organotrophic degradation are the Yin and Yang of the biological carbon cycle, involving a large number of organisms using a great diversity of metabolic pathways. Under oxic conditions, reduced monomeric compounds from biomass (e. g., carbohydrates, fatty acids.) are oxidized to carbon dioxide (CO_2_), which is then fixed again by autotrophic organisms to form reduced carbon compounds. Under anoxic conditions, e.g. in sediments of seas and lakes, reduced carbon compounds are only partly oxidized to carbon dioxide through fermentation and anaerobic respiration. In the absence of respiratory electron acceptors other than CO_2_ and methyl groups biomass is converted to methane and CO_2_. This multi-step process is brought about by a plethora of microorganisms belonging to different metabolic guilds, hydrolytic organisms, primary and secondary fermenting microorganisms, acetogenic bacteria and methanogenic archaea (methanogens) (1). First, polymeric carbon compounds are hydrolyzed to monomers (sugars, amino acids, nucleotides and fatty acids). These monomers are then fermented by primary fermenters to hydrogen (H_2_) and CO_2_, acetate and/or organic acids/alcohols (e. g., ethanol, propionate, butyrate, succinate). H_2_ and CO_2_ are both acetogenic and methanogenic substrates. Secondary fermenters utilize the organic acids and alcohols and convert them to H_2_, CO_2_ and acetate. The acetate produced can in turn be converted to CH_4_ and CO_2_ by aceticlastic methanogens (2). The low energy yield from methanogenesis forces the organisms to cooperate very efficiently with other organisms in the anaerobic food chain. While industrial application of this natural process, anaerobic digestion, produces biogas, the fragile interactions of multi-species food chains converting complex biomass to methane are easily disrupted, a major impediment to the efficient and reliable conversion of renewable biomass as an alternative to fossil fuels.

Compared to other groups of chemolithotrophic anaerobes (e. g., acetogenic and sulfate reducing bacteria), methanogens are metabolically restricted, both regarding the range of electron donors, and electron acceptors, they use. The former include only H_2_, C1 compounds, acetate, ethanol, and a few secondary alcohols (3). Unconventional electron donors for methanogenesis are, e. g., Fe^0^ (4), cathodes (5), carbon monoxide (6), and pyruvate (7, 8). It is curious that the latter is so far the most “complex” carbon compound methanogens can fully convert to CH_4_ and CO_2_. To the contrary, acetogenic bacteria are capable of funneling CO_2_ and reducing equivalents accruing from oxidation of all kinds of organic compounds into the reductive acetyl-CoA pathway (9).

The present study aimed at establishing glucose-dependent methanogenesis as a principle energy metabolism to support growth of *Methanosarcina acetivorans*. Its motivation was based on the following observations: (i) in natural environments lacking exogenous electron acceptors, this metabolic pathway occurs and requires the metabolic potential of at least two types of organisms, one acetogenic and one aceticlastic (10); (ii) *M. acetivorans* forms glycogen via gluconeogenesis and is capable of mobilizing the glycogen formed by converting it to methane (11); (iii) *M. acetivorans* is capable of growing methanogenically on pyruvate (8), which is a required link between glycolysis and the methanogenic energy metabolism in this organism; (iv) the only functions *M. acetivorans* appeared to lack for employing glucose-dependent methanogenesis were those for glucose-uptake and activation. Beside the principal value of the knowledge gained by experimentally expanding the metabolic capacity of *M. acetivorans* towards carbohydrate utilization, it would valorize the ever-increasing number of proposals deduced from metagenomic analyses, of methanogens using organic matter (12). Also, exploiting such advance into biotechnological applications would be an obvious benefit.

## Results

### Establishing glucokinase from *Escherichia coli* in *M. acetivorans*

Since *M. acetivorans* does not encode an obvious homolog of glucokinase involved in glycolysis (13), *glk* from *E. coli* was heterologously expressed in a tetracycline-(tet-) dependent fashion (by putting the structural gene under the control of the PmcrB(tetO1) promoter (14)). When encoded on a self-replicating episomal shuttle vector, Glk activity was up to two U mg^-1^ (Fig. 1A). When chromosomally integrated into the permissive *ssu* locus, which encodes a sulfonate transporter (strain CS1), tet-dependent Glk activity was much lower, approx. 70 mU mg^-1^, but still clearly above the background of approx. 5 mU mg^-1^(Fig. 1B).

**Fig. 1:**
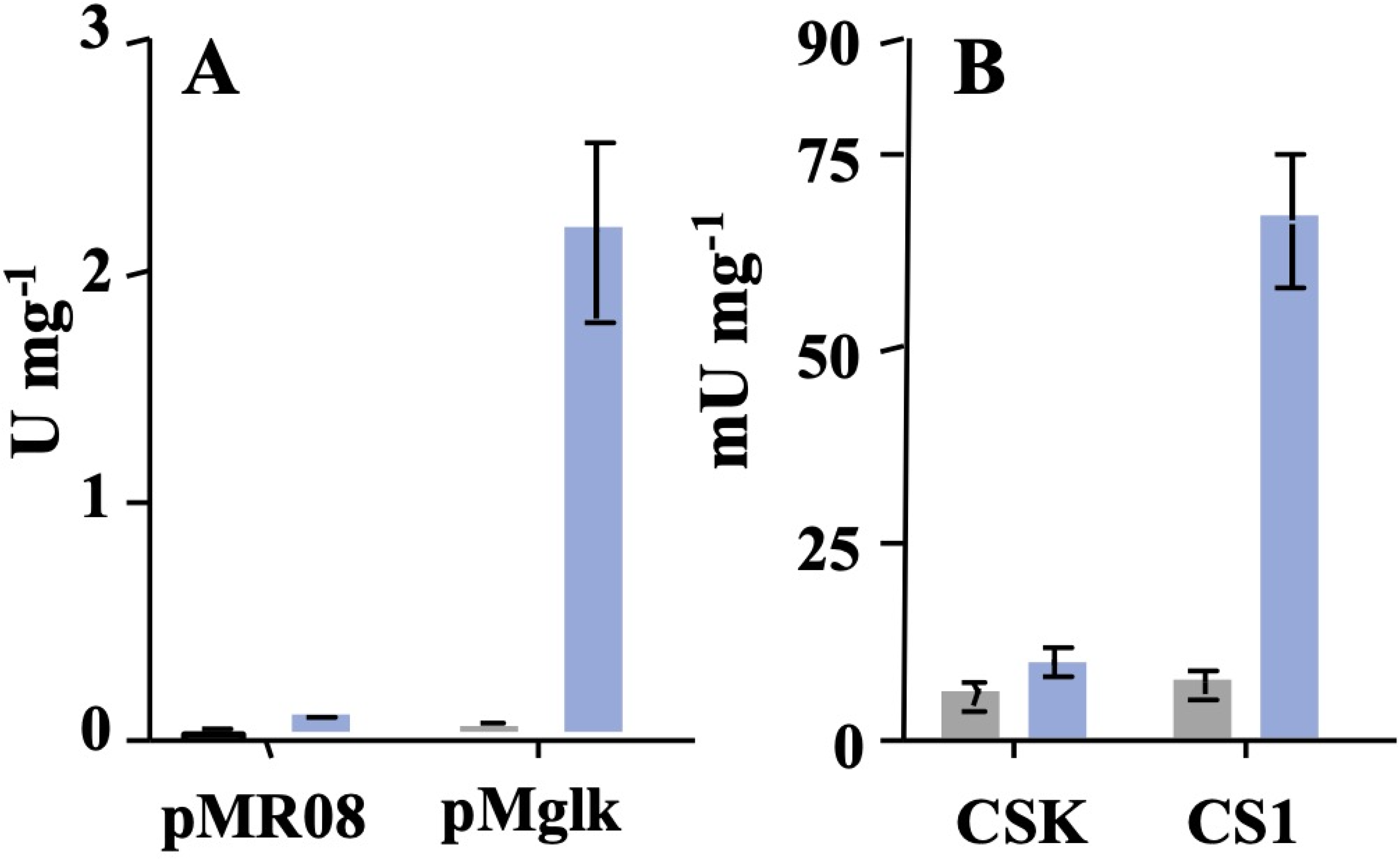
Heterologous glucokinase activity in *M. acetivorans*. Enzyme activity was quantified in cleared extracts of *M. acetivorans* strains. A: WWM73 with pMglk, *glk* in self-replicating vector. WWM73 with pMR08, control. B: CS1, *glk* chromosomally integrated (via pCSglk), CSK, isogenic control; bars represent the average of glucokinase activity in three biological replicates, the error bars designate the standard deviation; blue bars, cultivation with, grey bars, cultivation without tet; experiments were reproduced at least once.

### Establishing a glucose uptake facilitator in *M. acetivorans*

Since *M. acetivorans* does not encode any obvious function for carbohydrate uptake, such as a sugar phosphotransferase system, ATP binding cassette transporter, or a transporter for facilitated diffusion, *glf* from *Zymomonas mobilis* was heterologously expressed in a tet-dependent fashion. The structural gene was placed under the control of the PmcrB(tetO3) promoter, which leads to more than 17 times weaker expression than the PmcrB(tetO1) promoter (14). Our rationale was to tightly limit Glf synthesis, in order to minimize this bacterial protein to disturb integrity of the host’s archaeal membrane. A transporter for facilitated diffusion was chosen, based on the assumption, that (i) consumption of energy (as ATP or phosphoenolpyruvate) was not desirable in an energy-limited organism like *Methanosarcina*, and that (ii) intracellular glucose levels could be adjusted by the extracellular concentration using the Glf glucose facilitator. The *glf* gene was integrated into the chromosome of the *glk* expressing strain CS1, via replacing MA2965, which encodes a putative transposase (but lacking any vicinal transposon), resulting in strain CS3. Growth of CS3 on methanol was dramatically inhibited in the presence of glucose, and in a dose-dependent fashion (Fig. 2A and B). This “sensitivity” towards glucose was increased when tet was present (i. e., when *glk*/*glf* expression was induced) (Fig. 2B). Under this condition, already 0.1 mM glucose impaired methanol-dependent growth, whereas under non-inducing conditions 0.5 mM glucose was required for a similar effect, which indicates that the higher the *glk/glf* expression level, the more glucose “sensitivity” is attained in the transgenic organism. The fact that this “glucose phenotype” strictly depended on both Glf and glucose to be present provides solid evidence, albeit circumstantial, that the *glf trans*-gene is expressed and the glucose uptake facilitator (Glf) fully functional in the membrane of *M. acetivorans*. The effect of Glk/Glf in conjunction with glucose was also apparent in non-growing cells, where methanol-dependent methane formation was impaired by glucose (Fig. S1).

**Fig. 2:**
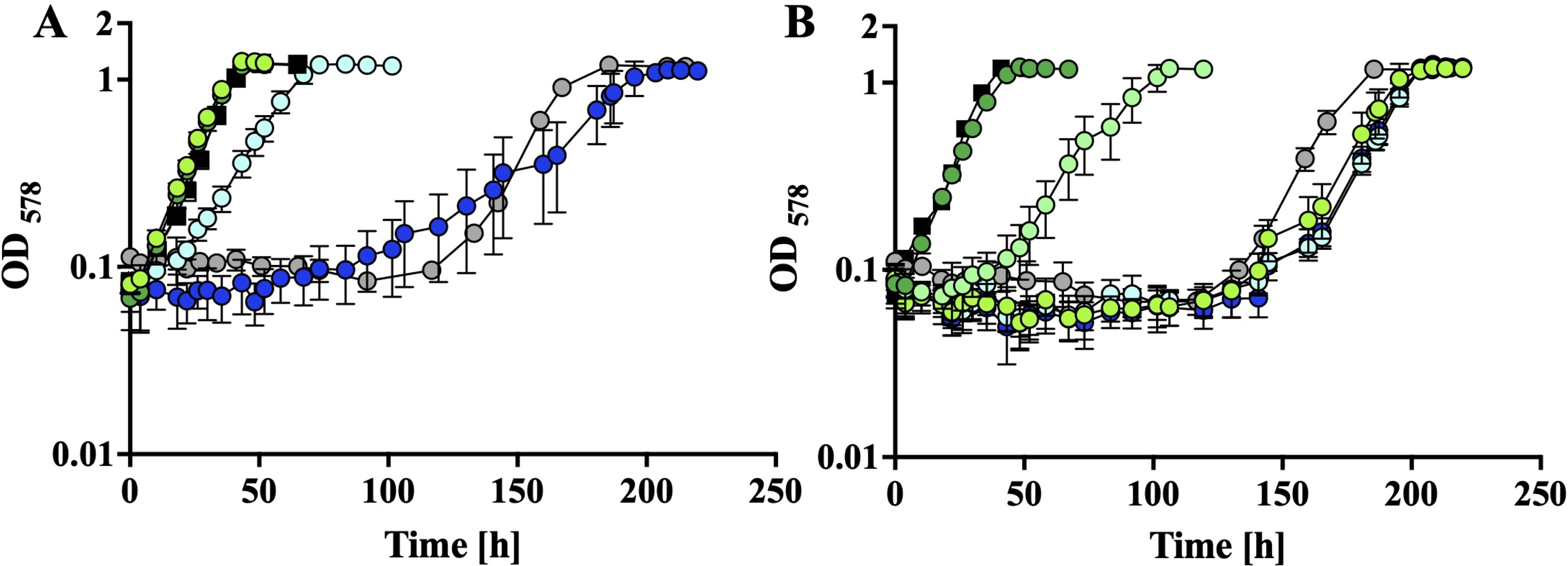
Glucose-dependent impairment of growth of *M. acetivorans* strains expressing *glk* and/or *glf*. Growth of strain CS3 on methanol without (A) and with (B) tet (100 µM) added, in the presence of no (dark green); 0.1 mM (turquoise), 0.25 mM (light green); 0.5 mM (cyan), 1 mM (dark blue), or 10 mM glucose (grey); as control, growth of strain CS1K on methanol, in the presence of 50 mM glucose (square) was included; shown are average values and their standard deviation as error bars of experiments with at least three biological replicates; experiments were reproduced at least once.

In the presence of more than 1.0 mM glucose, no growth on methanol could be observed, until after 90-120 h, when individual cultures started to grow at a similar rate and to a similar cell density than the control without glucose or the control without Glf (strain CS1K, with up to 50 mM glucose) (Fig. 2A and B). The observed glucose-dependent growth retardation of up to 150 h, abruptly followed by growth like the controls, was indicative of mutations arising in the population that suppress the glucose phenotype, either by eliminating *glf* expression, or by leading to inactive Glf. With the second transfer of the respective culture that arose in the presence of 1.0 mM glucose, it did not matter if glucose was present or not (Fig. S2A). Shifting that culture “back” to methanol-only for 10 transfers (approx. 35 generations), and subsequently to methanol plus 1.0 mM glucose, the suspected suppressor mutants again grew indistinguishably from CS3 without glucose (Fig. S2B), lending further support to the notion that (a) mutation(s) suppressing the glucose phenotype has/had occurred in CS3. Indeed, when four individual clones derived from this culture (obtained by single colony streak purification on methanol-containing media) were analyzed by sequencing PCR products containing *glf* (Tab. S1), three contained point mutations in the structural gene and one contained a point mutation at the position (+1) of transcription initiation in the (hybrid) promoter (Table S2). These four suppressor strains most likely have impaired *glf* expression or impaired Glf activity, although this notion was not investigated further.

We hypothesized that upon glucose uptake via Glf, glucose activation by the highly active Glk might be faster than the downstream reactions, effectively depleting the ATP pool of the cells. Alternatively, elevated levels of glucose-6-phosphate, the product of the reaction catalyzed by Glk, might be impairing normal cellular processes. Therefore, the strain expressing *glf* but lacking *glk*, CS3K (Table 2), was used for subsequent analyses. This strain also exhibited the glucose phenotype, but only at glucose concentrations higher than 1 mM (Fig. 3A). Since the glucose-dependent impairment of growth was again compounded by tet, it was omitted from subsequent experiments to avoid selection against *glf*. Strikingly, dose-dependent growth impairment by glucose coincides with a remarkable change in shape, size and light diffraction of CS3K upon exposure to growth-impairing amounts of glucose. At 1 mM, cells analyzed by phase contrast light microscopy appear larger, rounder, quasi “swollen”, and more light diffracting (Fig. 3B).

**Fig. 3:**
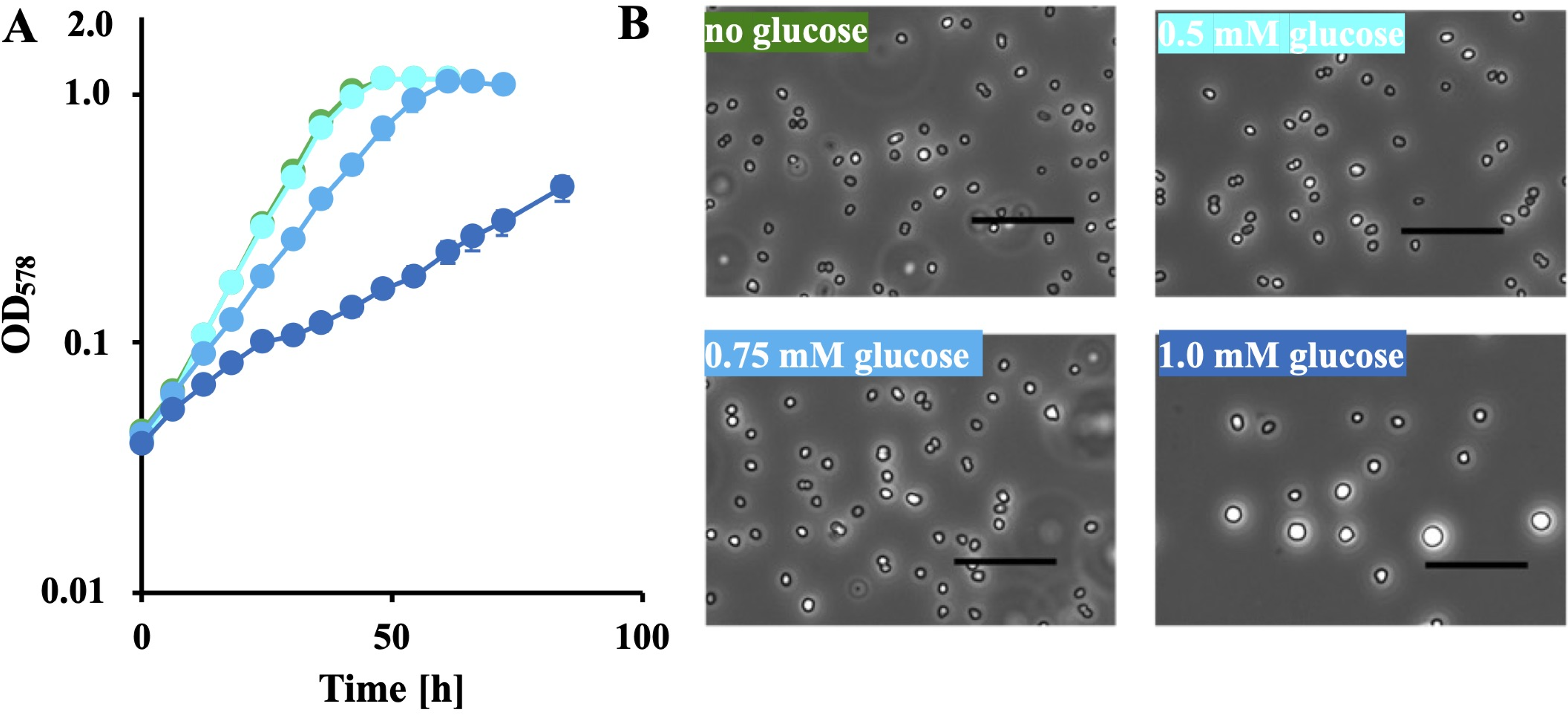
Glucose-dependent phenotype of *M. acetivorans* CS3K. A, growth of CS3K with methanol in the presence of 0.5 mM (cyan), 0.75 mM (light blue), 1 mM (blue) glucose and with methanol without glucose (green); shown are average values and their standard deviation as error bars of experiments with at least three biological replicates; B, phase contrast light micrographs of cells from (one replicate experiment of) A after 60 h; bars represent 20 µm.

### Tracing metabolism of exogenous glucose in *M. acetivorans*

While the growth phenotype and optical habitus indicated that the glucose was taken up and metabolized, it was unclear what to, considering the absence of Glk. To test if *M. acetivorans* could convert exogenous glucose to methane, experiments with non-growing cells (dense resting cell suspensions, see Materials and Methods) were conducted. In the presence of Glf, glucose was converted to methane at an initial rate of approx. 0.2 nmol min^-1^ mg^-1^, which was 10 times faster than without the transport function (approx. 0.02 nmol min^-1^ mg^-1^, Fig. 4A) but still 1,000 times slower than conversion of methanol (approx. 200 nmol min^-1^ mg^-1^, Fig. S1). When Glf was absent, glucose was not methanized, even when Glk was present (Fig. 4A). These experiments demonstrate that glycolytic methanogenesis could be established in *M. acetivorans*, and that this trait requires only Glf. Therefore, *M. acetivorans* must contain (an) endogenous function(s) for glucose activation. Next, glucose was used as the sole energy source in growth experiments to test if Glf-dependent methanogenesis from glucose could support growth of CS3K (Fig. 4B). In order to be able to follow both, gain and loss of biomass, higher than usual initial cell densities were chosen for this experiment and residual methanol was removed by washing the cells (see Materials and Methods). Without glucose, the optical density gradually declined, which is typical for *M. acetivorans* unable to grow. In the presence of 1 mM glucose, the optical density remained more or less constant, which could indicate that glucose can be used to provide (at least) the energy needed for “stationary” maintenance. However, given the observed change in shape, size and light diffraction, cell count (which was not determined in this experiment) could still have decreased, which makes the interpretation of this phenomenon difficult.

**Fig. 4:**
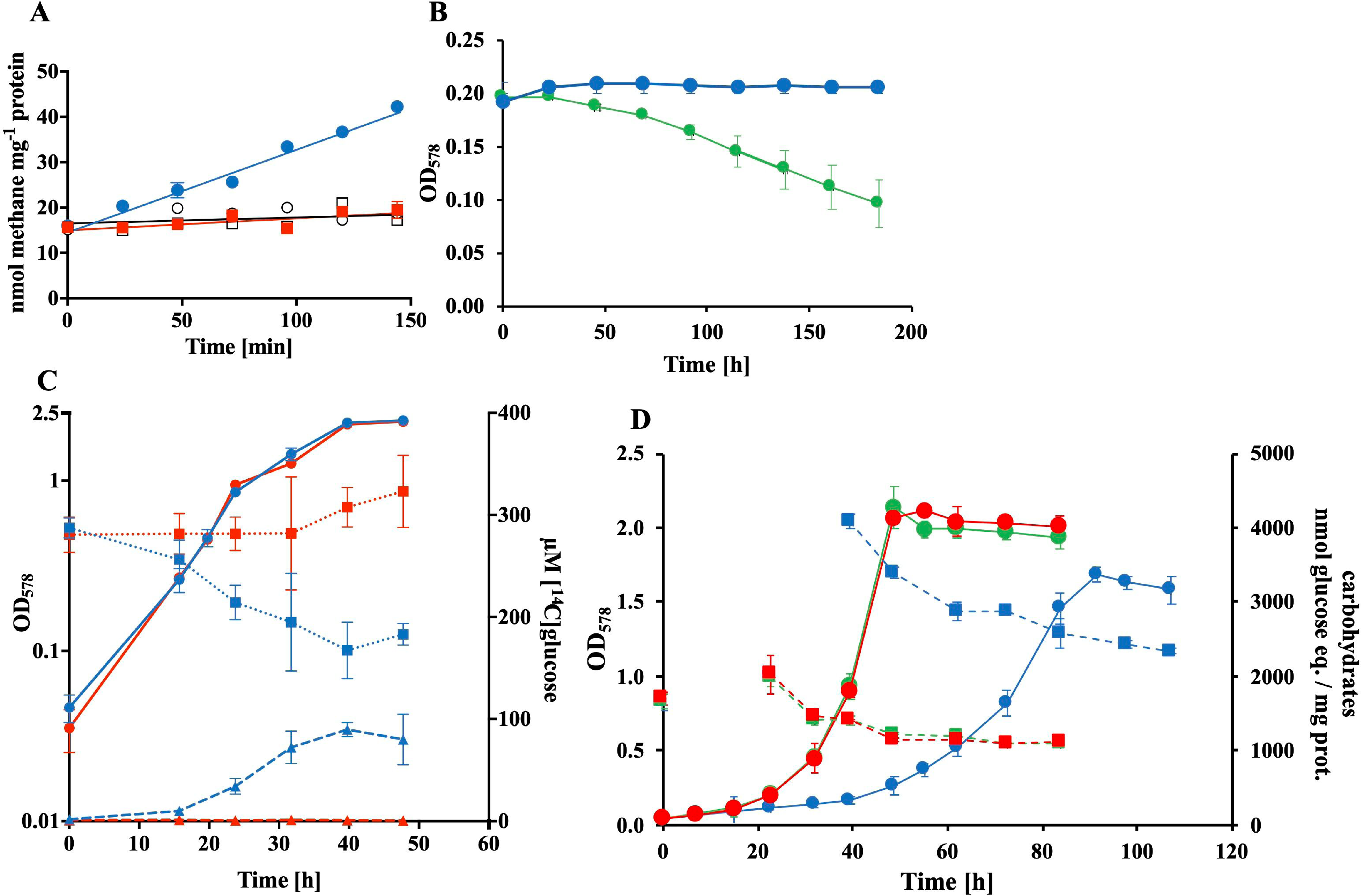
Conversion of glucose by *M. acetivorans*. A, methane formation in cell suspensions of CS3K (circles) and CS1K (squares), supplemented with 1 µmol glucose (filled symbols) or without glucose (open symbols); B, growth experiment using 1 mM glucose (blue) as sole energy source, or no energy source (green); C, conversion of [^14^C]glucose during growth (circles) by CS3K (blue) and CS1K (red) into non-gaseous metabolites; radioactivity in the culture supernatant (squares), and in the cell pellets (triangles); D, time course of intracellular accumulation of glucose and its derivative(s) (dashed lines) during growth (solid lines) on methanol of CS3K with no addition (green), or 1 mM glucose (blue) and of CSK-2 with 1 mM glucose (red); carbohydrate at time 0 h was determined in the precultures, the lack of data between 0 h and 23 h (red/green) and between 0 h and 40 h (blue) is due to insufficient accumulated biomass; shown are average values and their standard deviation as error bars of experiments with at least three biological replicates; experiments were reproduced at least once.

The slow rate of conversion in the cell suspension experiments and the inability to grow indicated that maybe only a fraction of exogenous glucose is converted to CH_4_ and CO_2_ by CS3K. In order to track the conversion of exogenous glucose into dissolved (i. e., non-gaseous and non-volatile) metabolites in growing cultures, uniformly ^14^C-labeled glucose (approx. 250 µM) was added to CS3K cultures growing on (non-radioactive) methanol, and glucose-derived metabolites were followed by determining the radioactivity in the culture supernatant and in the cells (culture sediments) (Fig. 4C). During the course of growth, a steady decrease in radioactivity could be observed in the supernatant, while radioactivity started to accumulate in the sedimented cells from 15 h onwards, again indicating glucose uptake in CS3K. No such uptake was observed in CS1K (Fig. 4C). Here, the amount of radioactivity in the supernatant remained almost constant over the course of the experiment, and there was no radioactivity detected in the sedimented cells. Notably, the total radioactivity resulting from the decrease of radioactivity in the supernatant and the increase of radioactivity in the cells after 40 h was close to the amount of radioactivity used in the experiment, i.e. the amount of glucose used. Intracellular glucose and its derivatives, like glycogen, were quantified in independent experiments (using unlabeled glucose) in cultures growing on methanol and with or without exogenous glucose added (Fig. 4D). In the presence of exogenous glucose, CS3K accumulates twice as much glucose-derived carbohydrate, than in its absence, or than a strain lacking Glf (Fig. 4D).

These experiments established, that glucose is taken up by and accumulating in CS3K, but is barely converted into gaseous and/or volatile metabolites. Together with the very slow glucose-dependent methane formation (Fig. 4A), glucose-dependent growth impairment (Fig. 3A) and the change in cell shape and size (Fig. 3B), some form of “metabolic block” appears to prevent further utilization of the “glucose” accumulated in the cells for energy conservation, such as glycolysis, followed by pyruvate-dependent methanogenesis (8). Apparently this “metabolic block” creates a strong negative selection leading to elimination of the heterologous Glf.

### Glucose as carbon source in *M. acetivorans*?

The observation that glucose-dependent methane formation is possible for strain CS3K (Fig. 4A) efforts were made to select via adaptive laboratory evolution (ALE) a strain growing on 10 mM glucose (i. e., conserve energy and synthesize biomass) by eliminating the assumed metabolic block through mutation. However, this approach was not successful even after three years of incubation. Given that already 1 mM glucose impaired methanol-dependent growth of *M. acetivorans* CS3K, efforts to select for a CS3K derivative able to co-metabolize methanol and glucose via ALE seemed futile, as no positive selection for this trait can be applied.

Instead, a *M. acetivorans* strain was created, the ability of which to grow should depend on its ability to utilize glucose anabolically, thus constituting a strong selection pressure for glucose utilization (instead of against). For this purpose, *glf* was inserted into strain MT28, a derivative of MCD21 which lacks both isoforms of the bifunctional carbon monoxide dehydrogenase/ acetyl-coenzyme A synthase (CODH/ACS) (15). Since the strain is not able to assimilate carbon from the energy substrate, methanol in this case, to acetyl-CoA, the strain’s energy and biosynthetic metabolism are “disconnected”, making addition of acetate necessary. We reasoned that with Glf, the strain could be “forced” to utilize glucose (instead of acetate) for its anabolic requirements and that the amount required would be much smaller than for catabolic purposes. The *glf*-containing *cdh* null mutant CS10 (Table 2), was shifted from medium containing methanol plus acetate to medium containing methanol plus 5 mM glucose (Fig. 5). Again, higher than usual initial cell densities were adjusted for this experiment and residual acetate in the preculture was removed by washing the cells (see Materials and Methods).

**Fig. 5:**
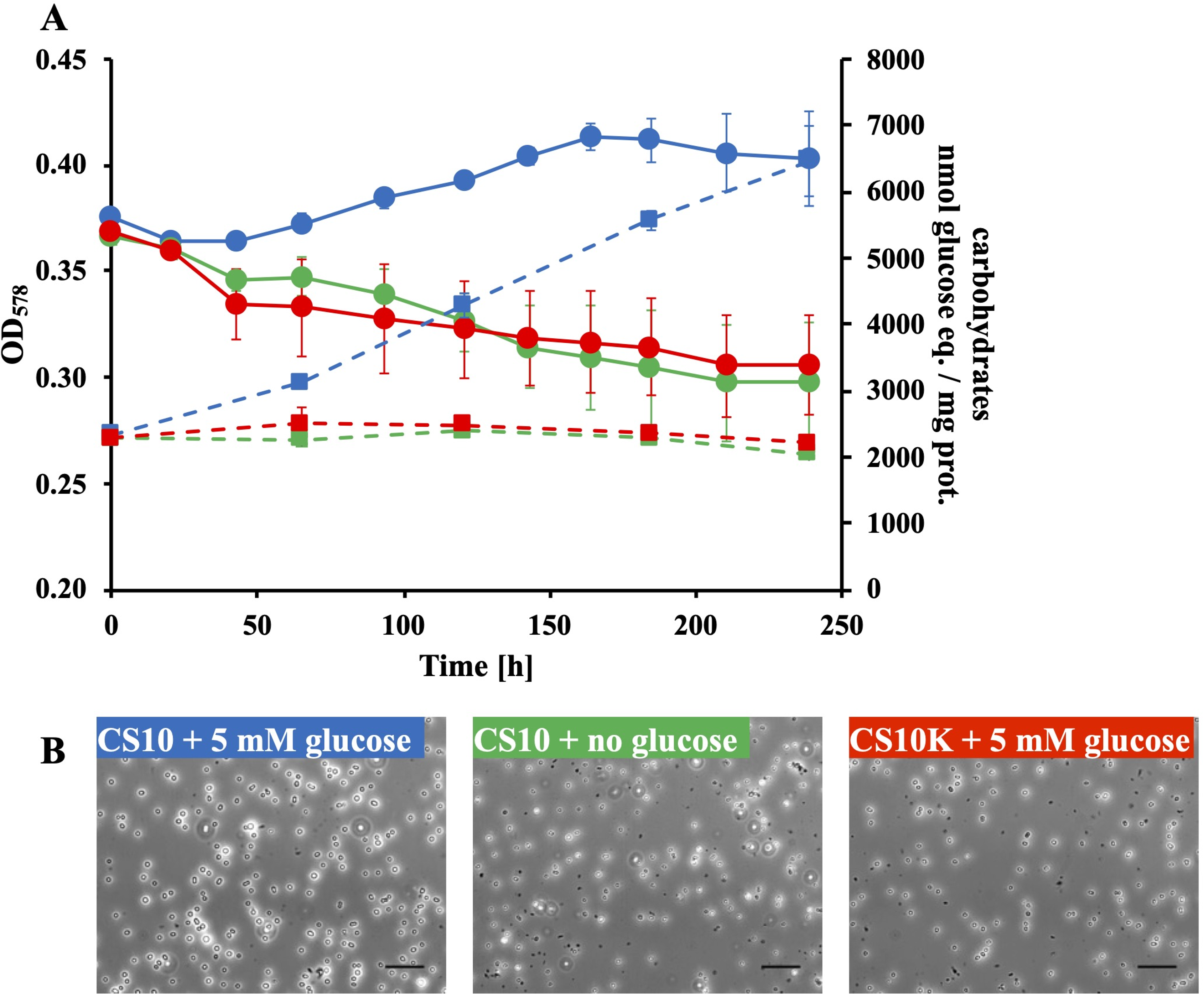
Glucose-dependent phenotype of *M. acetivorans* CS10. A, Growth of CS10 with methanol in presence of 5 mM glucose (blue) and with methanol without glucose (green) and CS10K (without Glf) with methanol in presence of 5 mM glucose (red); Carbohydrates were determined in the sedimented cells (dashed lines); shown are average values and their standard deviation as error bars of experiments with at least three biological replicates; B, phase contrast light micrographs of cells from (one replicate experiment of) A after 185 h.

Using glucose instead of acetate as the sole source for cell carbon in CS10 led to an increase of approx. 0.05 OD_578_ in the course of 120 h, while the values dopped for the same strain without glucose and the control strain (without *glf*) with glucose (Fig. 5A, solid lines). While these data again might indicate growth, and, thus, anabolic glucose utilization in CS10, change in cell shape, size and light refraction (Fig. 5B), previously observed in CS3K, makes this interpretation of the data unlikely. In fact, when transferred to new media after 250 h, no “growth” apparent from a change in OD_578_ could be observed for CS3K with methanol + glucose. Even after incubating the transferred cultures for more than two years no growth was apparent (except when contaminated, see discussion), indicating that ALE to select for anabolic glucose utilization is not possible under this condition. When the cultures were assessed for their carbohydrate content, CS10 accumulated approx. three times more in the presence of glucose than in its absence or than the control strain without Glf (Fig. 5A, dashed lines).

In order to more gently favor co-utilization of acetate and glucose for increased biomass production, i. e., to acclimatize CS10 to glucose without favoring loss of Glf, the strain was grown on methanol in the presence of limiting amounts of acetate (1 mM) and 0.2 mM glucose, which is not impairing growth (see also Fig. 2A) for more than 30 transfers (approx. 100 generations). However, omitting acetate led to the immediate cessation of growth.

## Discussion

While we clearly failed to construct a methanogen growing on glucose, valuable insights into *M. acetivorans*’s physiology and the physiological barriers for glucose utilization were gained.

### *M. acetivorans* can activate exogenous glucose

Expressing *glk* from *E. coli* proved to be counterproductive for establishing glucose-dependent methanogenesis as it compounded Glf- and glucose-dependent growth impairment. While we did not rigorously address this issue, it is reasonable to assume that the kinetic difference between ATP hydrolysis and ATP synthesis during glycolysis under this condition may have effectively depleted the ATP pool, which was reported to occur in other organisms (16). The residual glucokinase activity of approx. 5 mU mg^-1^ (Fig. 1B) falls in the range of other glycolytic activities determined in *M. acetivorans* (11) and is sufficient for activating exogenous glucose, once it is transported into the cell. The identity of this non-canonical enzyme is unknown but could be either a promiscuous kinase or a bifunctional phosphofructokinase (17), e. g., MA3563 (13).

### Functionality of Glf in *M. acetivorans*

Heterologous expression in *M. acetivorans* of *glf* from *Zymomonas mobilis*, which encodes a glucose uptake facilitator (18) led to impairment of methanol-dependent growth when glucose was present. Severeness of this phenotype depended on the concentration of glucose, consistent with the reported substrate affinity of the transporter (18). While again counterproductive for establishing glucose-dependent methanogenesis, this phenomenon provided indirect but conclusive evidence for the functionality of this bacterial transporter in *M. acetivorans*’s membrane. Since the glucose phenotype was observed in two unrelated genetic backgrounds (CS3K, CS10, Table 1), it is not a strain-specific but a *glf*-specific phenomenon in *M. acetivorans*.

**Table 1:**
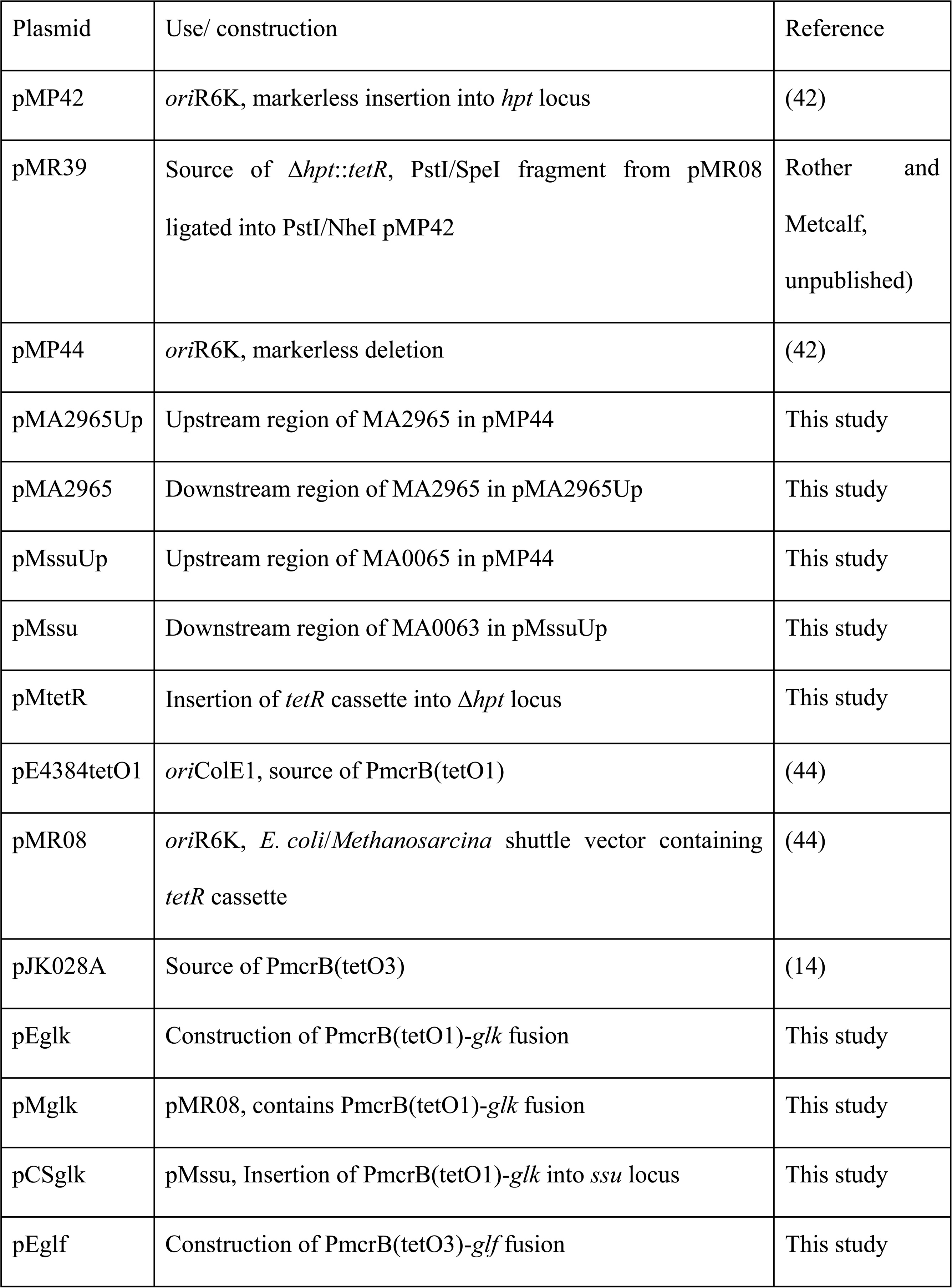
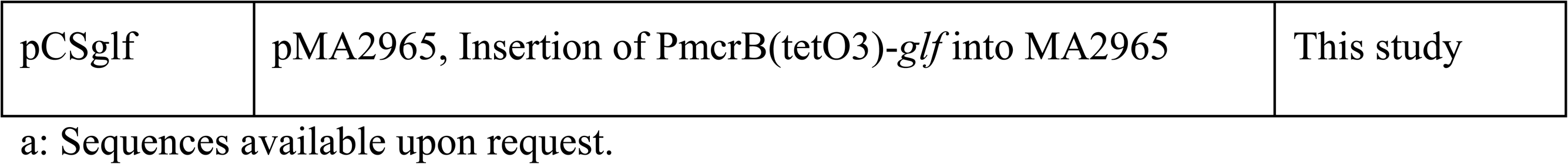
Plasmids used in this study^a^.

### “Glycogen” accumulation as reason or result of slow glycolysis?

Despite these caveats the transgenic *M. acetivorans* expressing *glf* was able to convert glucose to methane, but at a rate much slower than methanol was converted. Since methane formation from pyruvate is much faster in this organism (8), we conclude that the rate of glycolysis, not that of pyruvate conversion is limiting, which is expected as activities of the enzymes involved are typical for an “anabolic” pathway (11). Concomitant with glucose uptake, cells “swelled”, increased their light refraction, and accumulated (a) compound(s), which can be hydrolyzed to glucose. We did not investigate the identity of the carbohydrate-containing compound(s) the Glf^+^-strains accumulate, as *M. acetivorans* was shown to synthesize glycogen (11). It seems plausible that unphysiological accumulation of glucose, glucose-6-phosphate, or glycogen might impede canonical (methanol-dependent methanogenesis) and non-canonical (glucose-dependent methanogenesis) catabolism by altering the cell’s physico-chemical conditions, like water activity, charge distribution, or ATP/ADP-ratio. However, a metabolic block in the glycolytic pathway, or in a branching-off reaction involving a glycolytic intermediate could also slow the conversion of glucose, making accumulation of glycogen and the swelling/ light refraction change the consequence rather than the reason for the phenotype observed.

### Nature of the potential metabolic block insurmountable by ALE

We reasoned that the slow rate of glycolysis was causing ineffective glucose utilization. If increasing this rate was selectable, this trait should be evolvable, as *M. acetivorans* is remarkably susceptible for ALE (19–21). To make glucose utilization a positively selectable trait, *glf* was placed into an acetate auxotrophic *cdh* null mutant, which was then shifted from acetate to glucose as carbon source, both abruptly and gradually. In no case could a glucose-utilizing strain be selected, even after incubating for years. This result makes it highly unlikely that a regulatory effect is causing ineffective glycolysis. It was shown that *M. acetivorans* directs carbon through gluconeogenesis during exponential growth in order to accumulate storage glycogen, while changing the direction of carbon towards glycolysis during stationary growth phase in order to provide some maintenance energy (11). Such regulation should be easily overcome by ALE if survival of the organism depended on it. The same is true for the possible allosteric regulation of glycolytically active enzymes, e. g., inhibition of phosphofructokinase by ATP (22).

Instead, the selection phenotype observed suggests that either a toxic metabolite might be accruing during utilization of glucose, or a redox-imbalance. In neither case would it be easy to overcome the respective block by a single/a few spontaneous mutation(s). First, it was proposed that *Methanosarcina* species synthesize ribose-5-phosphate from triose phosphates via fructose-6-phosphate and involving hexulose-6-phosphate synthase releasing ribulose-5-phosphate and formaldehyde (23). Increasing metabolic flux through this pathway by increasing glycolysis (i. e., fructose-6-phosphate) might increase formaldehyde to a level the cells cannot readily detoxify (Fig. S3). Since ribose-5-phosphate and formaldehyde are stoichiometrically coupled and the former is a central anabolic metabolite, decreasing flux through the formaldehyde-generating path would be detrimental for the cells. Second, eliminating phosphate from glyceraldehyde-3-phosphate produces methylglyoxal (24), another toxic metabolite, yet required for biosynthesis of cofactor F_420_ (25). Again, increasing glycolysis might disturb the balance between making and using methylglyoxal (Fig. S3). Third, a potential bottleneck in connecting glycolysis to pyruvate-dependent methanogenesis could be the redox cofactors used. During glycolysis, *M. acetivorans* reduces NADP^+^ and/or NAD^+^ (11), during methanogenesis reduced ferredoxin, methanophenazine and F_420_ are used (26, 27). While electron transfer among those last three is well established, only reduced F_420_:NADP^+^ oxidoreductase (Fno) (28, 29), is known in methanogens. Whether electron exchange between NADH and F_420_ is possible in *M. acetivorans* is unknown, and lack of such enzyme would result in severe redox imbalance during glycolytic methanogenesis (Fig. S3).

### Biomass conversion to methane in a single organism?

All chemolithoautotrophic organisms require at least some gluconeogenetic aspects to their metabolism (e. g., for nucleotide biosynthesis) (30, 31). While many chemolithoautotrophic bacteria evolved (or acquired) heterotrophic traits, possibly to be able to compete in environments where organic compounds were abundant, methanogens retained their niches at the end of the anaerobic food chain (1). But why did they? One possible argument is the kinetic theory of optimal pathway length’s prediction that under planktonic conditions a glucose fermenting methanogen (Glucose → 3 CH_4_ + 3 CO_2_) would probably be less competitive than a syntrophic association of bacteria and archaea mediating the same reaction, despite the higher ATP yield (32). Heterotrophic organisms generally face a trade-off between rate and yield of ATP production, shorter pathways have a higher rate but lower yield (due to fewer coupling sites). This trade-off may result in an evolutionary dilemma, because cells with a higher rate but lower yield of ATP gain a selective advantage when competing for shared energy resources. Pathways with low rate and high yield, like methanogenic glucose utilization (Fig. S3), can be viewed as a form of cooperative resource use and may only evolve in spatially structured environments, like biofilms, which opens the possibility that glucose fermenting methanogens might once be found when isolated from biofilms. This leaves to explain why acetogens can easily switch from glucose fermentation to growth on H_2_ and CO_2_. Maybe the difference lies in the different electron carriers involved as has been discussed above.

### Direct glucose utilization for methanogenesis?

Considering all the caveats for glucose-dependent methanogenesis in *M. acetivorans* we encountered, a report containing data and conclusions, many of which grossly contradict the results and conclusions presented herein, came as a surprise (33). The most significant contradiction to the present data is the reportedly unimpaired growth of a *M. acetivorans* strain containing *glk* and *glf* genes (both from *Zymomonas mobilis*, both under the control of strong constitutive promoters) in the presence of 14 mM glucose (compare to Fig. 2 and Fig. 3) and the strain’s augmented methane yields compared to growth on methanol (33). Many experimental details are either not available (e. g., stability of *tran*s-genes, time of glucose exposure before assessment of growth physiology, assessment of culture purity, proportion of labelled to unlabeled metabolites, etc.) or differ in the two studies, which makes an in-depth comparison between the two neither meaningful nor appropriate. Still, two points are worth noting. First, we observed glucose-dependent augmentation of methanogenic growth only under conditions when contaminating microbes converted glucose to acetate, which could be used by *M. acetivorans* (Fig. S4). Second, if TCA intermediates were de novo synthesized in *M. acetivorans* from ^13^C glucose (i. e., from pyruvate and/or acetyl-CoA), completely different ^13^C incorporation than reported by (33) would result (24). Instead, the amount of ^13^C incorporated as reported by (33) is consistent with *M. acetivorans* using ^13^C acetate. Clearly, further studies are necessary to answer all open questions regarding glycolytic methanogenesis.

## Materials and Methods

### Strains and growth conditions

*Methanosarcina acetivorans* WWM73 and its derivatives (Table 2) were cultivated in high salt (HS) medium (34) containing trace elements and vitamins (35). Either 26-ml glass culture (Balch) tubes (Ochs, Bovenden, Germany) containing 10 ml, 100-ml serum bottles (Ochs) containing 20-50 ml, or 1300-ml infusion bottles (Ochs) containing 500 ml medium were used. Methanol (125 mM) (Sigma-Aldrich, Munich, Germany), supplemented from anaerobic, sterile stock, served as energy substrate. Acetate and/or glucose at the concentrations given for the respective experiments were added from anaerobic, sterile stock solutions. To remove residual methanol, cells were washed were indicated by twice centrifuging them anaerobically at 8,000 x g for 10 minutes, followed by resuspending them in substrate-free media. To avoid or remove bacterial contamination, glucose-containing cultures were occasionally supplemented with antibiotics (500 µg ml^-1^ ampicillin, 250 µg ml^-1^ kanamycin). Purity of cultures was confirmed via fluorescence microscopy (36) on a routine basis using an Axio Imager.M1 (Carl Zeiss Microscopy, Oberkochen, Germany) equipped with an HBO 100 light source and the shift free excitation filter set 18 (Carl Zeiss). Changes in morphology and light refraction of *M. acetivorans* strains carrying *glf* were documented by phase contrast microscopy using the same microscope, together with a AxioCam MRm camera and the AxioVision software (both Carl Zeiss).

**Table 2:**
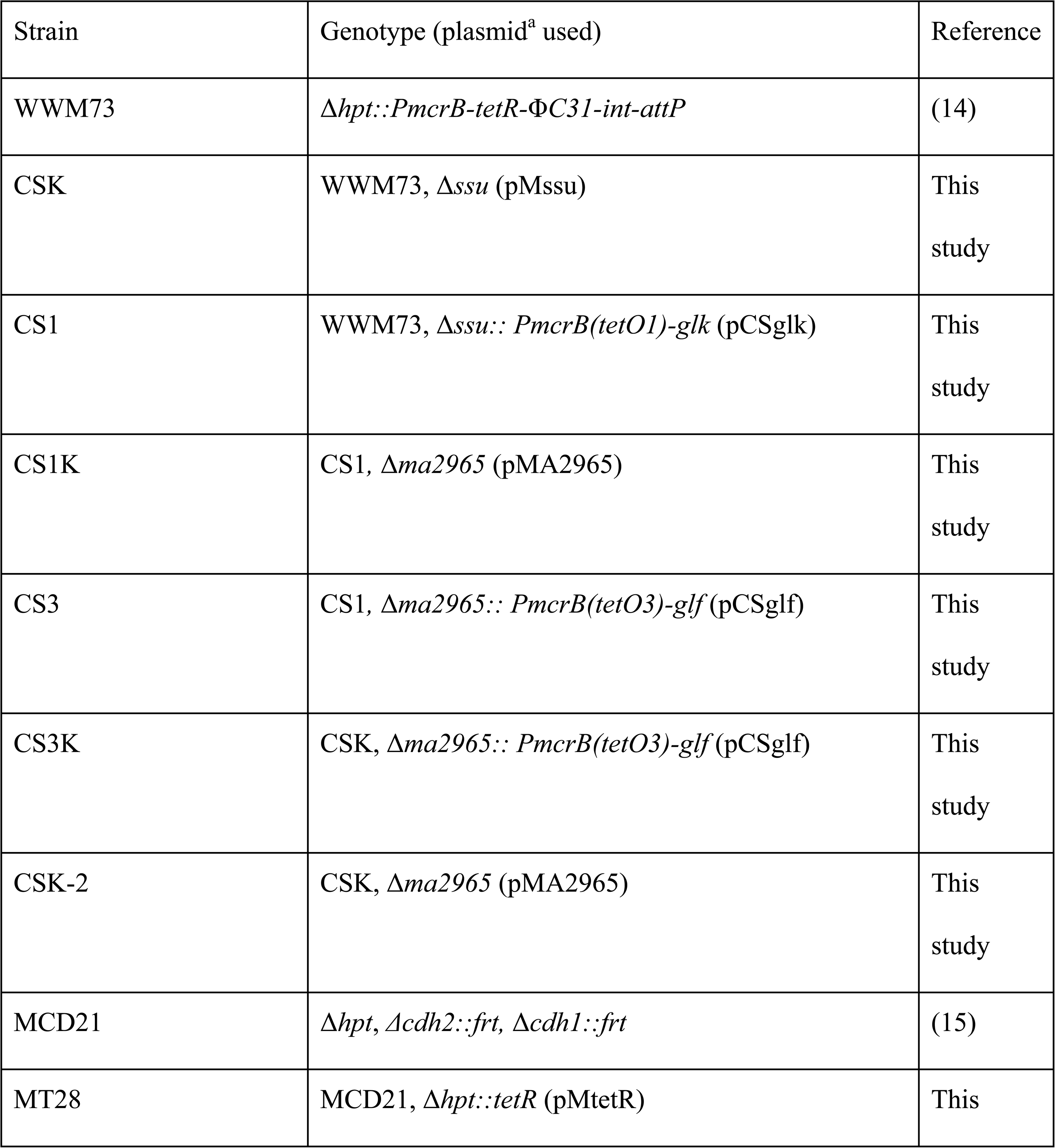
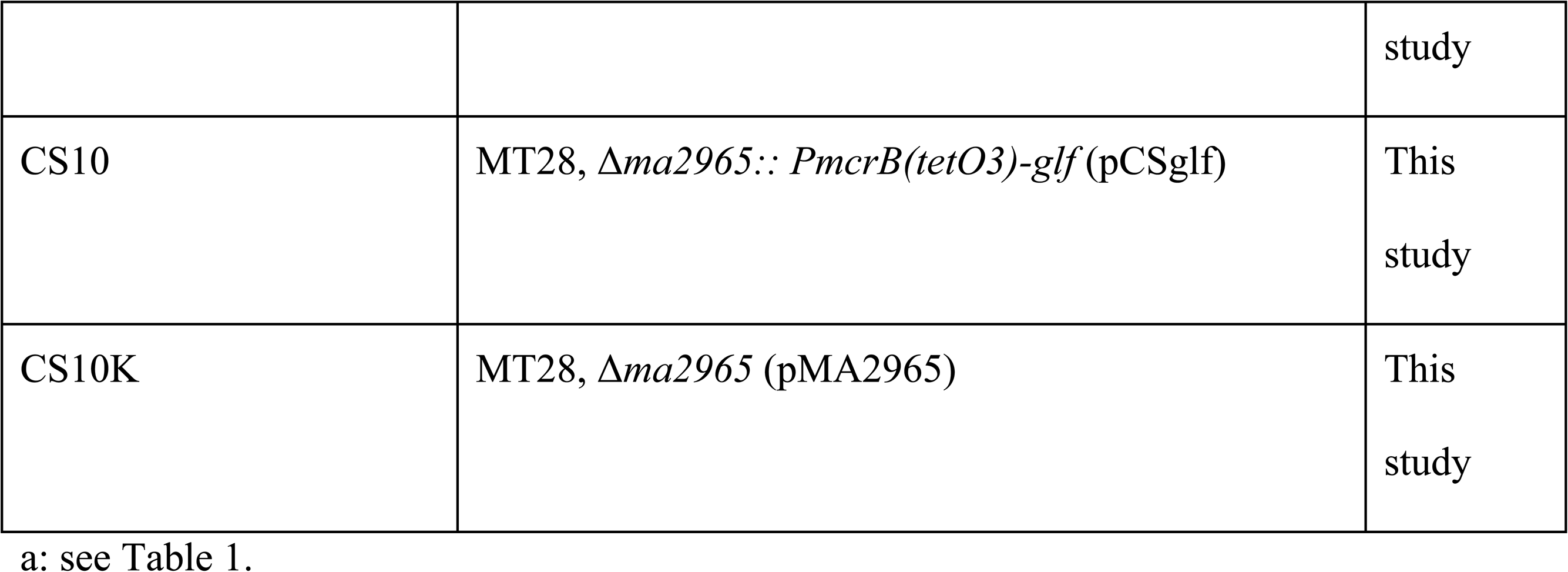
*M. acetivorans* strains used in this study.

Growth of *M. acetivorans* was monitored photometrically at 578 nm (OD_578_) either directly (i.e., undiluted) in Balch tubes using a Genesys 20 spectrophotometer (Thermo Scientific, Langenselbold, Germany) or using an Ultrospec 2000 spectrophotometer (Pharmacia Biotech, Uppsala, Sweden) for measurements of diluted culture samples.

### Molecular Methods and transformation

Standard conditions (37) were used for plasmid constructions and transformation of *Escherichia coli* DH10B (38) or WM1788 (for propagating R6K replicons) (39). Plasmids are presented in Table 1, oligonucleotides in Supplementary Table S1. Chromosomal DNA of microorganisms used in this study was isolated using a cetyl trimethylammonium bromide method (40) with modifications (34). All cloned PCR fragments were verified by Sanger sequencing using the BigDye Terminator Cycle Sequencing protocol (Microsynth Seqlab, Göttingen, Germany). Bacterial contamination of methanogenic cultures was assessed by 16S ribotyping, i. e., sequencing PCR fragments obtained using bacteria-specific oligonucleotides (Table S2) (41).

For construction of the plasmid pMssu, upstream and downstream regions (830 bp and 1,030 bp, respectively) of the *ssu* locus (MA0063-0065; annotated as sulfonate transporter) were amplified by PCR using the oligonucleotides listed in Supplementary Table S1 and chromosomal DNA from *M. acetivorans* WWM73 as template. The restriction sites AflII, PmlI, SphI and Eco32I were introduced into the upstream region and the sites KpnI, HpaI, NheI, NotI and BamHI into the downstream region. First, the DNA fragment of the upstream region was ligated into the plasmid pMP44 (42) via AflII and PmlI. The downstream DNA fragment was ligated into the resulting plasmid pMssuUp via HpaI and KpnI, resulting in the plasmid pMssu. This was either used to transform *M. acetivorans* for deleting the *ssu* locus, or a promoter-gene fusion construct was integrated between the upstream and downstream regions via SphI and NotI and subsequently used to transform *M. acetivorans*.

For construction of the plasmid pMA2965, upstream and downstream regions (1,020 bp and 1,070 bp, respectively) of the MA2965 locus (annotated as transposase gene) were amplified by PCR using the oligonucleotides listed in Supplementary Table S1 and chromosomal DNA from *M. acetivorans* WWM73 as template. The restriction sites AflII, PmlI, SphI and Eco32I were introduced into the upstream region and the sites KpnI, HpaI, NheI, NotI and BamHI into the downstream region. First, the DNA fragment of the upstream region was ligated into the plasmid pMP44 (42) using the restriction sites AflII and PmlI. The downstream DNA fragment was ligated into the resulting plasmid pMA2965Up (Table 1) via HpaI and KpnI, resulting in pMA2965 (Table 1). This plasmid was either used to transform *M. acetivorans* in order to eliminate the MA2965 locus, or a promoter-gene fusion construct was integrated between the upstream and downstream regions via SphI and NotI and subsequently used to transform *M. acetivorans*.

The plasmid pMR39 (Table 1) contains the *tetR* cassette (encoding the tetracycline-dependent regulator (43)) flanked by the upstream and downstream regions of the *hpt* locus of *M. acetivorans*. This region was excised using AflII and BamHI and blunted with Klenow fragment (Fisher Scientific, Schwerte, Germany). The fragment was then cloned into the vector pMP44 (42) via PmlI. The resulting plasmid was designated pMtetR and used to insert the *tetR* cassette into the Δ*hpt* locus of *M. acetivorans* strain MCD21 (15).

To allow regulated expression of genes in *M. acetivorans*, they were fused to a tetracycline-regulatable hybrid promoter via NdeI to PmcrB(tetO1) or PmcrB(tetO3) (14) and subcloned into pE4384tetO1 via SphI and NotI. Subsequently, the promoter gene fusions were either moved into the self-replicating plasmid pMR08 (44) or into the integration plasmids pMssu or pMA2965 (see above) via SphI and NotI. PmcrB(tetO1) was amplified by PCR from pE4384tetO1 (44) using oligonucleotides 1+2 and restricted with SphI and NdeI in order to fuse it with the structural genes via the NdeI site. The PmcrB(tetO3) was amplified by PCR from pJK028A (14) using the same oligonucleotides and proceeding analogously to PmcrB(tetO1).

The *glk* gene was amplified by PCR using the oligonucleotides listed in Supplementary Table S1 and chromosomal DNA from *E. coli* as template. It was fused to PmcrB(tetO1) giving rise to pEglk (Table 1). The fusion was moved to pMR08 and pMssu, resulting in pMglk and pCSglk (Table 1), respectively. The *glf* gene was amplified by PCR using the oligonucleotides listed in Supplementary Table S1 and chromosomal DNA from *Zymomonas mobilis* subsp. *mobilis* (DSM424), obtained from the Deutsche Sammlung von Mikroorganismen und Zellkulturen (Braunschweig, Germany), as template. It was fused to PmcrB(tetO3) giving rise to pEglf (Table 1). The PmcrB(tetO3)-*glf* fusion was moved to pMA2965 resulting in pCSglf (Table 1).

The plasmids were used to genetically modify *M. acetivorans* as described (44–46). Markerless deletion of *ssu* or MA2965 with or without inserting a *trans*-gene was conducted as described for *hpt* (42). The strains used in this study are presented in Table 2. The relevant genotype of the *M. acetivorans* strains constructed was verified via Southern hybridization using a digoxigenin system (Roche) as described (15).

### Cell suspension experiments

For determining initial rates of glucose-dependent methane formation, cell suspension experiments were conducted. All manipulations were carried out under strictly anaerobic conditions inside an anaerobic glove box (Coy, Grass Lake, USA) containing N_2_:H_2_ (96:4, v/v) or by using anaerobic and gas-tight containers. Cells were grown on methanol to late exponential growth phase, harvested by centrifugation and washed three times using an anaerobic buffer [50 mM Piperazine-*N*,*N*’-bis-(2-ethansulfonate), 478 mM NaCl, 13 mM KCl, 2 mM MgCl_2_, 2 mM CaCl_2_, 2.8 mM cysteine-HCl, 0.4 mM Na_2_S and 0.4 µM resazurin, pH 7.0 (NaOH)]. Cells were resuspended in 10 ml of the same buffer in a Balch tube and kept on ice while protein concentration in whole cells was determined as described (47) using bovine serum albumin as standard. Cells were diluted in anaerobic buffer to a protein concentration of 2-4 mg ml^-1^ and a final volume of 5 ml. Puromycin (2 µg ml^-1^) was added from a sterile anaerobic stock solution to prevent de novo protein biosynthesis. After changing the gas atmosphere to 50 kPa N_2_ suspensions were incubated in a shaking water bath at 37 °C for 20-24 h in order to deplete the cells of storage compounds, which could be converted to methane (11), and, thus, to obtain a stable baseline for measuring very slow methane formation. Subsequently, the gas phase was again changed to 50 kPa N_2_. Measurements were initiated by adding either 100 mM methanol or 1 mM glucose. Initial methane formation rates were determined of the course of 2.5 h.

### Metabolite analyses

Methane in the gas phase was quantified using a GC-2010 Plus gas chromatograph (Shimadzu GmbH, Diusburg, Germany) equipped with an Optima-5 column (length, 30 m; diameter, 0.25 µm; film thickness, 0.25 µm) (Macherey-Nagel, Düren, Germany) developed with N_2_ as carrier gas at a column flow rate of 1.7 ml min^-1^. The temperature of the flame ionization detector was set to 200 °C, that of the column oven to 130 °C, and that of the split (20:1) injector to 280 °C. Gas phase samples (50 µl) were injected with a gas-tight sample lock syringe (Hamilton, Bonaduz, Switzerland).

Glucose in supernatants of cultures was quantified enzymatically using the glucose (GO) assay kit (Sigma-Aldrich, St. Lois, USA), according the manufacturer’s instructions and calibrating with a standard curve using authentic *D*-(+)-glucose dissolved/diluted like the respective analytes.

Carbohydrate accumulated in *M. acetivorans* (“glycogen”) was quantified using a phenol/sulfuric acid method (48). Briefly, cultures or cell suspensions were harvested by centrifugation (8,000 x g. 10 min at 8 °C) and pellets containing up to 1.6 mg whole cell protein (47) were lysed in deionized water containing 0.1 µg ml^-1^ RNase A (Roth, Karlsruhe, Germany) and 0.1 µg ml^-1^ DNase I (Roche, Mannheim, Germany), respectively. The lysate was cleared by centrifugation at 13,000 x g for 10 minutes and the supernatant constituted the cleared cell-free lysate. Of this, 0.4 ml (appropriately diluted in deionized water, if required) was mixed with 0.2 ml 5% (v/v) Roti-Aqua-Phenol (Roth, Karlsruhe, Germany) before 1 ml sulfuric acid [< 95 % (v/v) VWR International, Darmstadt, Germany] was added. The samples were vigorously mixed, incubated for 20 minutes at 30 °C in thermomixer at 300 rpm (Eppendorf, Hamburg, Germany) and the extinction at 490 nm (against water) recorded using an Ultrospec 2000 spectrophotometer (Pharmacia Biotech, Uppsala, Sweden). For calibration, glycogen from bovine liver (Sigma) was used, dissolved and treated identically as the analyte samples.

### Enzyme activity

Glucokinase (Glk) activity was determined using cleared cell-free lysates (see above) of *M. acetivorans* at room temperature by coupling the reaction to the formation of NADPH by glucose-6-phosphate dehydrogenase (49). A reaction mixture of 1 ml contained 50 mM Tris-HCl, pH 8.0; 13.3 mM MgCl_2_, 112 mM glucose, 550 µM ATP, 230 µM NADP^+^ and 10 U glucose-6-phosphate dehydrogenase (Roche, Mannheim, Germany). The enzymatic reaction was started with the addition of 50 µl cell-free lysate and NADPH formation was monitored over 270 s at 340 nm. Specific Glk activity was calculated using the molar extinction coefficient for NADPH of ε= 6.22 x 10^3^ l/mol^-1^ x cm^-1^ (50) and the protein concentration in the cleared cell-free lysate, determined according to (51) and using bovine serum albumin as standard, and is given in U (1U = 1 µmol NADP^+^ reduced) per mg protein.

## Acknowledgments

This work was supported by the Center for Synthetic Microbiology (SYNMIKRO), Marburg, Germany, to M.Ro.. The funders had no role in study design, data collection and interpretation, or the decision to submit the work for publication. We are grateful to Rudolf K. Thauer (Marburg) for his support, for fruitful discussions, and critical comments on this manuscript.

## Author contribution

M.Ro. conceived and supervised the study, C.Sa., M.Ri., and M.Ro. designed experiments, C.Sa. and M.Ri. acquired the data, all authors analyzed and interpreted the data, M.Ro. drafted the manuscript, all authors revised the manuscript.

## References

1. Schink B. 1997. Energetics of syntrophic cooperation in methanogenic degradation. Microbiol Mol Biol Rev 61:262–280.

2. Ferry JG. 1992. Methane from acetate. J Bacteriol 174:5489–5495.

3. Whitman WB, Bowen TL, Boone DR. 2006. The Methanogenic Bacteria, p 165–207. *In* Dworkin M, Falkow S, Rosenberg E, Schleifer K-H, Stackebrandt E (ed), The Prokaryotes-A Handbook on the Biology of Bacteria, 3rd ed, vol 3. Springer, New York.

4. Uchiyama T, Ito K, Mori K, Tsurumaru H, Harayama S. 2010. Iron-corroding methanogen isolated from a crude-oil storage tank. Appl Environ Microbiol 76:1783–1788.

5. Lohner ST, Deutzmann JS, Logan BE, Leigh J, Spormann AM. 2014. Hydrogenase-independent uptake and metabolism of electrons by the archaeon *Methanococcus maripaludis*. ISME J 8:1673–1681.

6. Rother M, Metcalf WW. 2004. Anaerobic growth of *Methanosarcina acetivorans* C2A on carbon monoxide: An unusual way of life for a methanogenic archaeon. Proc Natl Acad Sci USA 101:16929–16934.

7. Bock AK, Priegerkraft A, Schönheit P. 1994. Pyruvate - a novel substrate for growth and methane formation in *Methanosarcina barkeri*. Arch Microbiol 161:33–46.

8. Richter M, Sattler C, Schöne C, Rother M. 2024. Pyruvate-dependent growth of *Methanosarcina acetivorans*. J Bacteriol 206:e00363–00323.

9. Schuchmann K, Müller V. 2016. Energetics and application of heterotrophy in acetogenic bacteria. Appl Environ Microbiol 82:4056–4069.

10. Winter J, Wolfe RS. 1979. Complete degradation of carbohydrate to carbon dioxide and methane by syntrophic cultures of *Acetobacterium woodii* and *Methanosarcina barkeri*. Arch Microbiol 121:97–102.

11. Santiago-Martinez MG, Encalada R, Lira-Silva E, Pineda E, Gallardo-Perez JC, Reyes-Garcia MA, Saavedra E, Moreno-Sanchez R, Marin-Hernandez A, Jasso-Chavez R. 2016. The nutritional status of *Methanosarcina acetivorans* regulates glycogen metabolism and gluconeogenesis and glycolysis fluxes. FEBS J 283:1979–1999.

12. Chadwick GL, Skennerton CT, Laso-Pérez R, Leu AO, Speth DR, Yu H, Morgan-Lang C, Hatzenpichler R, Goudeau D, Malmstrom R, Brazelton WJ, Woyke T, Hallam SJ, Tyson GW, Wegener G, Boetius A, Orphan VJ. 2022. Comparative genomics reveals electron transfer and syntrophic mechanisms differentiating methanotrophic and methanogenic archaea. PLoS Biol 20:e3001508.

13. Galagan JE, Nusbaum C, Roy A, Endrizzi MG, Macdonald P, FitzHugh W, Calvo S, Engels R, Smirnov S, Atnoor D, Brown A, Allen N, Naylor J, Stange-Thomann N, DeArellano K, Johnson R, Linton L, McEwan P, McKernan K, Talamas J, Tirrell A, Ye W, Zimmer A, Barber RD, Cann I, Graham DE, Grahame DA, Guss AM, Hedderich R, Ingram-Smith C, Kuettner HC, Krzycki JA, Leigh JA, Li W, Liu J, Mukhopadhyay B, Reeve JN, Smith K, Springer TA, Umayam LA, White O, White RH, Conway de Macario E, Ferry JG, Jarrell KF, Jing H, Macario AJ, Paulsen I, Pritchett M, Sowers KR, Swanson RV, Zinder SH, Lander E, Metcalf WW, Birren B. 2002. The genome of *M. acetivorans* reveals extensive metabolic and physiological diversity. Genome Res 12:532–542.

14. Guss AM, Rother M, Zhang JK, Kulkarni G, Metcalf WW. 2008. New methods for tightly regulated gene expression and highly efficient chromosomal integration of cloned genes for *Methanosarcina* species. Archaea 2:193–203.

15. Matschiavelli N, Oelgeschläger E, Cocchiararo B, Finke J, Rother M. 2012. Function and regulation of isoforms of carbon monoxide dehydrogenase/acetyl-CoA synthase in *Methanosarcina acetivorans*. J Bacteriol 194:5377–5387.

16. Teusink B, Walsh MC, van Dam K, Westerhoff HV. 1998. The danger of metabolic pathways with turbo design. Trends Biochem Sci 23:162–169.

17. Zamora RA, Gonzalez-Ordenes F, Castro-Fernandez V, Guixe V. 2017. ADP-dependent phosphofructokinases from the archaeal order *Methanosarcinales* display redundant glucokinase activity. Arch Biochem Biophys 633:85–92.

18. Parker C, Barnell WO, Snoep JL, Ingram LO, Conway T. 1995. Characterization of the *Zymomonas mobilis* glucose facilitator gene product (*glf*) in recombinant *Escherichia coli*: examination of transport mechanism, kinetics and the role of glucokinase in glucose transport. Mol Microbiol 15:795–802.

19. Rother M, Boccazzi P, Bose A, Pritchett MA, Metcalf WW. 2005. Methanol-dependent gene expression demonstrates that methyl-CoM reductase is essential in *Methanosarcina acetivorans* C2A and allows isolation of mutants with defects in regulation of the methanol utilization pathway. J Bacteriol 187:5552–5559.

20. Schöne C, Poehlein A, Jehmlich N, Adlung N, Daniel R, von Bergen M, Scheller S, Rother M. 2022. Deconstructing *Methanosarcina acetivorans* into an acetogenic archaeon. Proc Natl Acad Sci USA 119:e2113853119.

21. Bao J, Somvanshi T, Tian Y, Laird M, Garcia P, Schöne C, Rother M, Borrel G, Scheller S. 2024. Nature AND Nurture: Enabling formate-dependent growth in *Methanosarcina acetivorans*. bioRxiv doi:10.1101/2024.01.08.574737.

22. Verhees CH, Tuininga JE, Kengen SW, Stams AJ, van der Oost J, de Vos WM. 2001. ADP-dependent phosphofructokinases in mesophilic and thermophilic methanogenic archaea. J Bacteriol 183:7145–7153.

23. Goenrich M, Thauer RK, Yurimoto H, Kato N. 2005. Formaldehyde activating enzyme (Fae) and hexulose-6-phosphate synthase (Hps) in *Methanosarcina barkeri*: a possible function in ribose-5-phosphate biosynthesis. Arch Microbiol 184:41–48.

24. Grochowski LL, White RH. 2008. New and unconventional metabolism in methanogenic Archaea. Ann N Y Acad Sci 1125:190–214.

25. Grochowski LL, Xu H, White RH. 2006. Identification of lactaldehyde dehydrogenase in *Methanocaldococcus jannaschii* and its involvement in production of lactate for F_420_ biosynthesis. J Bacteriol 188:2836–2844.

26. Deppenmeier U, Müller V. 2008. Life close to the thermodynamic limit: how methanogenic archaea conserve energy. Results Probl Cell Differ 45:123–152.

27. Thauer RK, Kaster AK, Seedorf H, Buckel W, Hedderich R. 2008. Methanogenic archaea: ecologically relevant differences in energy conservation. Nat Rev Microbiol 6:579–591.

28. Yamazaki S, Tsai L. 1980. Purification and properties of 8-hydroxy-5-deazaflavin-dependent NADP^+^ reductase from *Methanococcus vannielii*. J Biol Chem 255:6462–6465.

29. Berk H, Thauer RK. 1998. F_420_H_2_:NADP oxidoreductase from *Methanobacterium thermoautotrophicum*: identification of the encoding gene via functional overexpression in *Escherichia coli*. FEBS Lett 438:124–126.

30. Ronimus RS, Morgan HW. 2003. Distribution and phylogenies of enzymes of the Embden-Meyerhof-Parnas pathway from archaea and hyperthermophilic bacteria support a gluconeogenic origin of metabolism. Archaea 1:199–221.

31. Fuchs G. 2011. Alternative pathways of carbon dioxide fixation: insights into the early evolution of life? Annu Rev Microbiol 65:631–658.

32. Costa E, Perez J, Kreft JU. 2006. Why is metabolic labour divided in nitrification? Trends Microbiol 14:213–219.

33. Ma JY, Yan Z, Sun XD, Jiang YQ, Duan JL, Feng LJ, Zhu FP, Liu XY, Xia PF, Yuan XZ. 2024. A hybrid photocatalytic system enables direct glucose utilization for methanogenesis. Proc Natl Acad Sci U S A 121:e2317058121.

34. Metcalf WW, Zhang JK, Shi X, Wolfe RS. 1996. Molecular, genetic, and biochemical characterization of the *serC* gene of *Methanosarcina barkeri* Fusaro. J Bacteriol 178:5797–5802.

35. Sowers KR, Noll KM. 1995. Techniques for anaerobic growth, p 15-47. *In* Sowers KR, Schreier HJ (ed), Methanogens, vol 2. Cold Spring Harbor Laboratory Press, Plainview, NY.

36. Doddema HJ, Vogels GD. 1978. Improved identification of methanogenic bacteria by fluorescence microscopy. Appl Environ Microbiol 36:752–754.

37. Ausubel FM, Brent R, Kingston RE, Moore DD, Seidmann JG, Smith JA, Struhl K (ed). 2003. Curr. Protoc. Mol. Biol. J. Wiley & Sons, Inc., New York.

38. Grant SG, Jessee J, Bloom FR, Hanahan D. 1990. Differential plasmid rescue from transgenic mouse DNAs into *Escherichia coli* methylation-restriction mutants. Proc Natl Acad Sci U S A 87:4645–4649.

39. Haldimann A, Wanner BL. 2001. Conditional-replication, integration, excision, and retrieval plasmid-host systems for gene structure-function studies of bacteria. J Bacteriol 183:6384–6393.

40. Murray MG, Thompson WF. 1980. Rapid isolation of high molecular weight plant DNA. Nucleic Acids Res 8:4321–4325.

41. DeLong EF. 1992. Archaea in coastal marine environments. Proc Natl Acad Sci U S A 89:5685–5689.

42. Pritchett MA, Zhang JK, Metcalf WW. 2004. Development of a markerless genetic exchange method for *Methanosarcina acetivorans* C2A and its use in construction of new genetic tools for methanogenic archaea. Appl Environ Microbiol 70:1425–1433.

43. Beck CF, Mutzel R, Barbe J, Müller W. 1982. A multifunctional gene (*tetR*) controls Tn*10*-encoded tetracycline resistance. J Bacteriol 150:633–642.

44. Oelgeschläger E, Rother M. 2009. *In vivo* role of three fused corrinoid/methyl transfer proteins in *Methanosarcina acetivorans*. Mol Microbiol 72:1260–1272.

45. Metcalf WW, Zhang JK, Apolinario E, Sowers KR, Wolfe RS. 1997. A genetic system for Archaea of the genus *Methanosarcina*: liposome-mediated transformation and construction of shuttle vectors. Proceedings of the National Academy of Sciences of the United States of America 94:2626–2631.

46. Boccazzi P, Zhang JK, Metcalf WW. 2000. Generation of dominant selectable markers for resistance to pseudomonic acid by cloning and mutagenesis of the *ileS* gene from the archaeon *Methanosarcina barkeri* Fusaro. J Bacteriol 182:2611–2618.

47. Schmidt K, Liaanen Jensen S, Schlegel HG. 1963. Die Carotinoide der *Thiorodaceae*. Arch Mikrobiol 46:117–126.

48. DuBois M, Gilles KA, Hamilton JK, Rebers PA, Smith F. 1956. Colorimetric Method for Determination of Sugars and Related Substances. Anal Chem 28:350–356.

49. Fraenkel DG, Horecker BL. 1964. Pathways of D-glucose metabolism in *Salmonella typhimurium*: A study of a mutant lacking phosphoglucose isomerase. J Biol Chem 239:2765–2771.

50. Bergmeyer HU (ed). 1974. Methoden der Enzymatischen Analyse. Verlag Chemie Weinheim/Bergstrasse.

51. Bradford MM. 1976. A rapid and sensitive method for the quantitation of microgram quantities of protein utilizing the principle of protein-dye binding. Anal Biochem 72:248–254.

